# Sex differences and estradiol effects in MAPK and Akt cell signalling across subregions of the hippocampus

**DOI:** 10.1101/2021.05.30.446341

**Authors:** Paul A. S. Sheppard, Tanvi A. Puri, Liisa A. M. Galea

**Author notes:** Address correspondence to: Dr. Liisa Galea, Djavad Mowafaghian Centre for Brain Health, University of British Columbia, 2215 Wesbrook Mall, Vancouver, BC, Canada, V6T 1Z3. Email addresses: PASS –, TAP –, LAMG –. Schulich School of Medicine and Dentistry, Robarts Research Institute, University of Western Ontario, London, ON, Canada.

## Abstract

**Introduction:** Rapid effects of estrogens within the hippocampus of rodents are dependent upon cell signaling cascades, and activation of these cascades by estrogens varies by sex. Whether these pathways are rapidly activated within the dentate gyrus (DG) and CA1 by estrogens and across the anatomical longitudinal axis has been overlooked.

**Methods:** Gonadally-intact female and male rats were given either vehicle or physiological systemic low (1.1µg/kg) or high (37.3µg/kg) doses of 17β-estradiol thirty minutes prior to tissue collection. To control for the effects of circulating estrogens, an additional group of female rats was ovariectomized (OVX) and administered 17β-estradiol. Brains were extracted and tissue punches of the CA1 and DG were taken along the longitudinal hippocampal axis (dorsal and ventral) and analyzed for key MAPK and Akt cascade phosphoproteins.

**Results:** Intact females had higher Akt pathway phosphoproteins (pAkt, pGSK-3β, pp70S6K) than males in the DG (dorsal, ventral) and lower pERK1/2 in the dorsal DG. Most effects of 17β-estradiol on cell signalling occurred in OVX animals. In OVX animals, 17β-estradiol increased cell signalling of MAPK and Akt phosphoproteins (pERK1/2, pJNK, pAkt, pGSK-3β) in the CA1 and pERK1/2 and pJNK DG.

**Discussion/Conclusions:** Systemic 17β-estradiol treatment rapidly alters phosphoprotein levels in the hippocampus dependent on reproductive status and intact females have greater expression of Akt phosphoproteins than intact males across the hippocampus. These findings shed light on underlying mechanisms of sex differences in hippocampal function and response to interventions that affect MAPK or Akt signaling.

## Introduction

Estrogens rapidly affect object and spatial memory consolidation (1–3), short-term social memory (2,4), response learning (5), neurogenesis (6), and spine and synapse formation (7–9). Most of these effects of estrogens occur rapidly and likely via cell signaling pathways after binding with the estrogen receptors (ERs). Estrogens exert both rapid (non-classical) and delayed (classical) actions. The classical actions of estrogens occur when they bind with ERs that dimerize, translocate to the nucleus, and act directly to affect gene expression (10). In contrast, the rapid effects of estrogens initiate a myriad of intracellular processes, including the activation of cell signaling cascades (reviewed in (2–4)). Rapid (within minutes) or delayed (within hours) effects of estrogens on molecular, cellular, and behavior can vary by brain region (reviewed in (2,3,11)). Rapid effects of estrogens on social, object, and spatial location recognition rely on signaling through mitogen-activated protein kinase (MAPK) and Akt (protein kinase B) pathways, whereas the delayed effects are driven predominantly by ER-estrogen response element interactions and effects on gene expression (2,3). There is, however, overlap in these effects, with the less well understood rapid effects occurring concurrently with the more established delayed effects. Thus, understanding the mechanisms, such as cell signaling cascades, through which the rapid effects act is imperative to a comprehensive view of actions by estrogens.

Two of the most studied cell signaling cascades are the MAPK and Akt pathways. These pathways are involved in a number of processes including cell proliferation, migration, and survival, apoptosis, metabolism, differentiation, immune responses, and development (12–15). There is much crosstalk between MAPK and Akt pathways (16,17), and these pathways often act in parallel to elicit similar downstream mechanisms and outputs in complementary or synergistic manners (16,18,19). However, little work has investigated both pathways in the same animals and those that do typically only measure specific phosphoproteins of interest, despite the intricacy and interconnectedness of these signaling cascades.

Sex differences have been noted in the influence of 17β-estradiol on the hippocampus. In older rats, acute 17β -estradiol increases spine density in older females but decreases spine density in males (20). Furthermore, whereas gonadectomy reduces spine density in both males and females in the CA1 (21–23), estradiol increases spine density in females (24,25) but not in males (22). Few studies have investigated sex differences in the effects of 17β-estradiol on cell signalling. Female rats have higher baseline levels of phosphorylated extracellular signal-regulate protein kinase 1/2 (pERK1/2; both p44 [ERK1] and p42 [ERK2] isoforms), pAkt, and phosphorylated glycogen synthase kinase 3β (pGSK-3β in the prefrontal cortex (prelimbic and infralimbic cortices), nucleus accumbens, and rostral caudate putamen of rats than male rats (26). However, to our knowledge, no studies have examined sex differences in expression of both MAPK and Akt phosphoproteins in different subregions of the hippocampus thus far. Few studies have explored sex differences in these cell signalling proteins after exposure to 17β-estradiol. Intrahippocampal 17β-estradiol increases dorsal hippocampal pAkt and pERK2 in OVX females, but not in gonadally-intact or castrated males (27). Similarly, 17β-estradiol increases pERK1/2 in anteroventral periventricular (AVPV) nucleus and the medial preoptic area in females but not in males (28). Thus, there is a dearth of evidence exploring influence of 17β-estradiol on cell signalling across the sexes and within subregions across the hippocampus.

The hippocampus is a heterogeneous structure with distinct subfields across the longitudinal axis (dorsal, ventral), as well as dentate gyrus (DG), CA3 and CA1 subregions. Based on gene expression, receptor levels, activity, and connectivity, the dorsal hippocampus is thought to be important for learning and memory, whereas the ventral region is more important for regulation of stress and anxiety (29). Estradiol rapidly increases cell proliferation in both the dorsal and ventral DG (6,30). Furthermore, 17β-estradiol increases apical dendritic spines in the CA1 region but not in the DG (21,31). Given that 17β-estradiol influences both dorsal and ventral hippocampal plasticity, it is important to examine possible subregional (i.e. DG and CA1) effects across the longitudinal axis that show rapid 17β-estradiol action in both sexes.

Most previous studies have examined the effects of estrogens on cell signaling in either the whole hippocampus (32), dorsal hippocampus (27,33–35), or dorsal CA1 (36) in OVX female rodents. Estrogens rapidly increase phosphorylated (activated) proteins in the MAPK and Akt cascades including the MAPKs pERK and pJNK and Akt pathway proteins pAkt, pGSK-3β, and p70S6K. Another MAPK protein, p38, has divergent effects from other MAPKs and is minimally affected in the hippocampus by the delayed effects of 17β-estradiol treatment in OVX female rats (37). However, even in studies investigating the rapid effects of 17β-estradiol on cell signaling cascades, methodologies (including, for example, timing and/or dose of 17β-estradiol treatment) vary and make comparison between pathways and proteins difficult.

The current study was designed to examine sex differences in cell signaling cascades, with and without 17β-estradiol, in the dorsal and ventral fields of DG and CA1 of the hippocampus. We hypothesized that sex and 17β-estradiol would differentially affect MAPK and Akt cell signaling cascades and that these differences would vary by region across the longitudinal axis. Furthermore, we hypothesized that in OVX rats 17β-estradiol would increase activation of cell signaling cascades in both the CA1 and DG, but that these effects may be more prominent in the dorsal versus ventral axis.

## Methods

### Animals

Ten-to eleven-week-old, gonadally intact female (n=65) and male (n=19) Sprague-Dawley rats (Charles River, St. Constant, Quebec, Canada) were received at the University of British Columbia and pair-housed (with the exception of one triple-housed cage of males) upon arrival. Rats were housed in transparent polyurethane bins (48×27×20cm) with aspen chip bedding and were given Purina rat chow and tap water *ad libitum*. Rats were maintained under a 12/12h light/dark cycle (lights on 07:00h) in standard housing conditions (21 ± 1°C; 50 ± 10% humidity). Animals were handled for 5 minutes each, every other day prior to the experiment for a minimum of five handling sessions. Animals were 12-13 weeks old when administered 17β-estradiol and brains were collected. The means and standard errors of the means for body mass were as follows: females 295±3.55g and males 477±7.51g. To determine estrous phase during testing, intact females underwent vaginal lavage immediately following tissue collection. Lavage samples were stained with cresyl violet and estrous phase was evaluated (38). One week following arrival, 36 female rats were bilaterally ovariectomized using aseptic techniques under isoflurane anesthesia, with ketamine (30 mg/kg, Bimeda-MTC, Cambridge, ON), xylazine (2 mg/kg, Bayer HealthCare, Toronto, ON), and bupivacaine (applied locally; 4 mg/kg, Hospira Healthcare Corporation, Montreal, QC). Following surgery, rats were single-housed and allowed to recover for at least one week before testing. All experiments were conducted in accordance with the ethical guidelines set by the Canada Council for Animal Care and were approved by the University of British Columbia Animal Care Committee. All efforts were made to reduce the number and the suffering of animals.

### Treatment

Rats were subcutaneously administered 1.1µg/kg 17β-estradiol in sesame oil (low 17β-estradiol; Females n=10, Males n=6, OVX n=13), 37.3µg/kg 17β-estradiol in sesame oil (high 17β-estradiol; Females n=10, Males n=7, OVX n=13), or sesame oil vehicle (Females n=9, Males n=6, OVX n=10) 30 minutes prior to tissue extraction. These sample sizes were selected as they have previously produced large effect sizes (Cohen’s d>0.8) in similar analyses (6,39). Estradiol doses were chosen because they result in circulating levels of 17β-estradiol observed on the morning of proestrus (high 17β-estradiol) or during diestrus (low 17β-estradiol)(40), and both doses enhance cell proliferation 30 minutes after injection in OVX females (6,30). There were no significant differences in average body mass per treatment group (all ps>0.12). All rats were injected between 09:00h and 11:15h. Rats were sacrificed by decapitation, and brains were excised, hemisected, and flash-frozen on dry ice.

### Brain tissue processing

Brains were sliced into 300µm sections at -10°C using a Leica CM3050 S cryostat (Nußloch, Germany). Punches were taken from the DG and CA1 of both the dorsal and ventral horns of the hippocampus using modified 18G needles with an internal punch diameter of 0.838mm (see Figure 1). Tissue was collected and homogenized using an Omni Bead Ruptor (Omni international, Kennesaw, GA) with 40μl of cold lysis buffer. Homogenates were centrifuged at 4°C and 1000×g for 15min then stored at -20°C. Total protein concentrations in homogenates were quantified using the Pierce Micro BCA Protein Assay Kit (ThermoFisher Scientific) and used according to manufacturer instructions, with samples run in triplicates.

**Figure 1:**
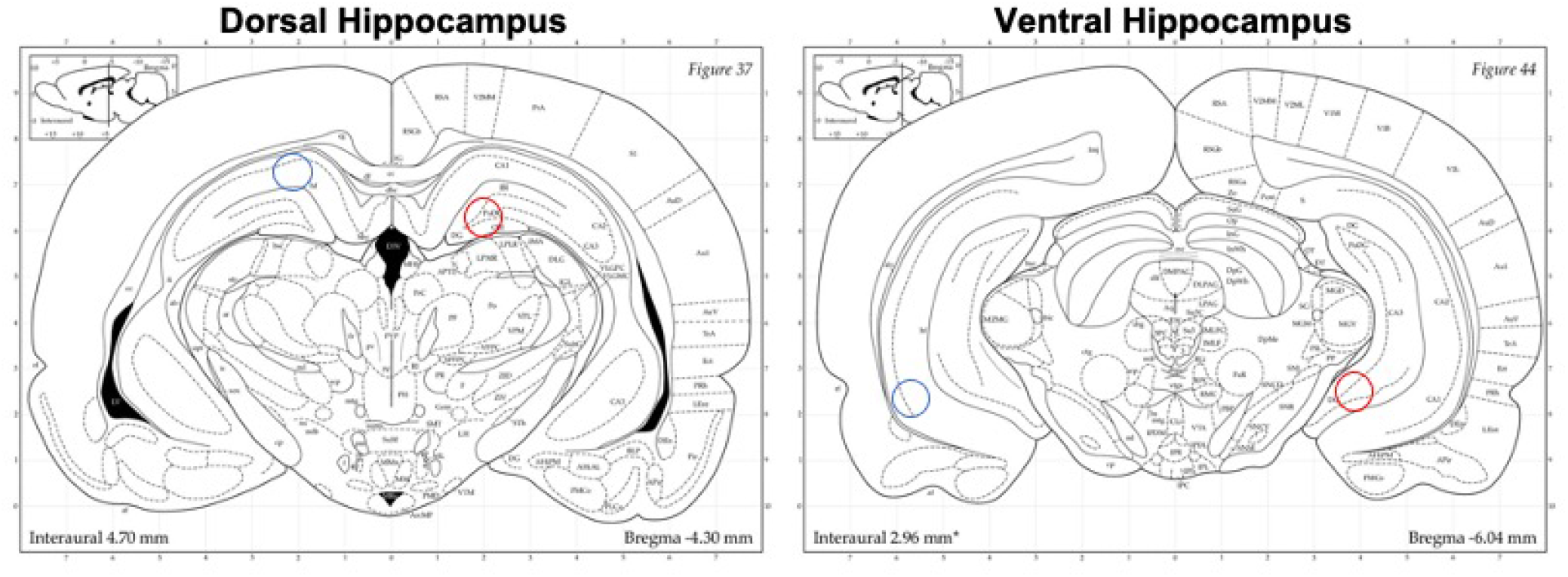
Example of CA1 and dentate gyrus punch location. Blue circles indicate approximate CA1 tissue punch location. Red circles indicate approximate dentate gyrus tissue punch location. Punches were 0.838mm in diameter.

### Phosphoprotein quantification

Multiplex electrochemiluminescence immunoassay kits (MAPK Kinase Phosphoprotein and Akt Signaling Panel Assay Whole Cell Lysate Kits) were purchased from Meso Scale Discovery (Rockville, MD) and used according to manufacturer instructions – with the exception of incubating the samples at 4°C overnight – to measure phosphoprotein levels. Electrochemiluminescent assays are highly selective and quantitative, produce low background, have a broad dynamic range of detection, and require relatively small volume of sample compared to ELISAs and Western blots (41,42). The antibody precoated plates allowed for the simultaneous quantification of the following phosphoproteins: pERK1/2 (combined p42 and p44 isoforms), pJNK, and pp38 (MAPK kit) and pAkt, pGSK-3β, and pp70S6K (Akt kit). Samples were run in technical replicates and plates were read with a Sector Imager 2400 (Meso Scale Discovery) using the Discovery Workbench 4.0 (Meso Scale Discovery). Phosphoprotein values were averaged across technical replicates and normalized to total protein concentrations for statistical analyses (as in (39,43–45)). Inter-and intra-assay coefficients of variation (CV) were below 10%. Phosphoprotein signal data were excluded when the CV between duplicates exceeded this threshold.

### Statistical analyses

Statistical analyses were run using Statistica v.8.0 (StatSoft Inc, Tulsa, OK) software. Bartlett’s homogeneity of variance tests were performed and, in analyses where homogeneity of variance was violated, data were subjected to a Box-Cox transformation (lambda=-0.5). Two-way ANOVAs were conducted for each analyte and region with treatment (oil, low 17β-estradiol, high 17β-estradiol) and sex (female, male) as between groups factors. To determine if there were baseline differences between males and females, vehicle treated rats were used as reference for calculation of percent differences. Phosphoprotein levels in intact vehicle treated female rats were compared across the estrous cycle (proestrus vs non-proestrus, corresponding to high and low 17β-estradiol phases, respectively) using Welch’s t-tests. One-way ANOVAs were conducted for each analyte and region in the OVX animals, with treatment (vehicle, low 17β-estradiol, high 17β-estradiol) as the between-subjects factor. *Post hoc* comparisons used Newman-Keuls. When one-way ANOVAs violated homogeneity of variance, Brown-Forsythe ANOVAs were performed with Dunnett’s T3 *post hocs. A priori* analyses were subjected to Bonferroni correction. Any statistical outliers (± 2 standard deviations from the group mean) were removed. Statistical significance was set at p<0.05. Eta squared (η^2^) and Cohen’s d effect sizes were calculated where appropriate.

## Results

First, we analysed the effects of estrous cycle on phosphoproteins across the dorsal/ventral axis and in all regions. There were no significant effects (all ps>0.245; Supplementary Figures 1 and 2) and, as such, these were not considered further.

**Figure 2:**
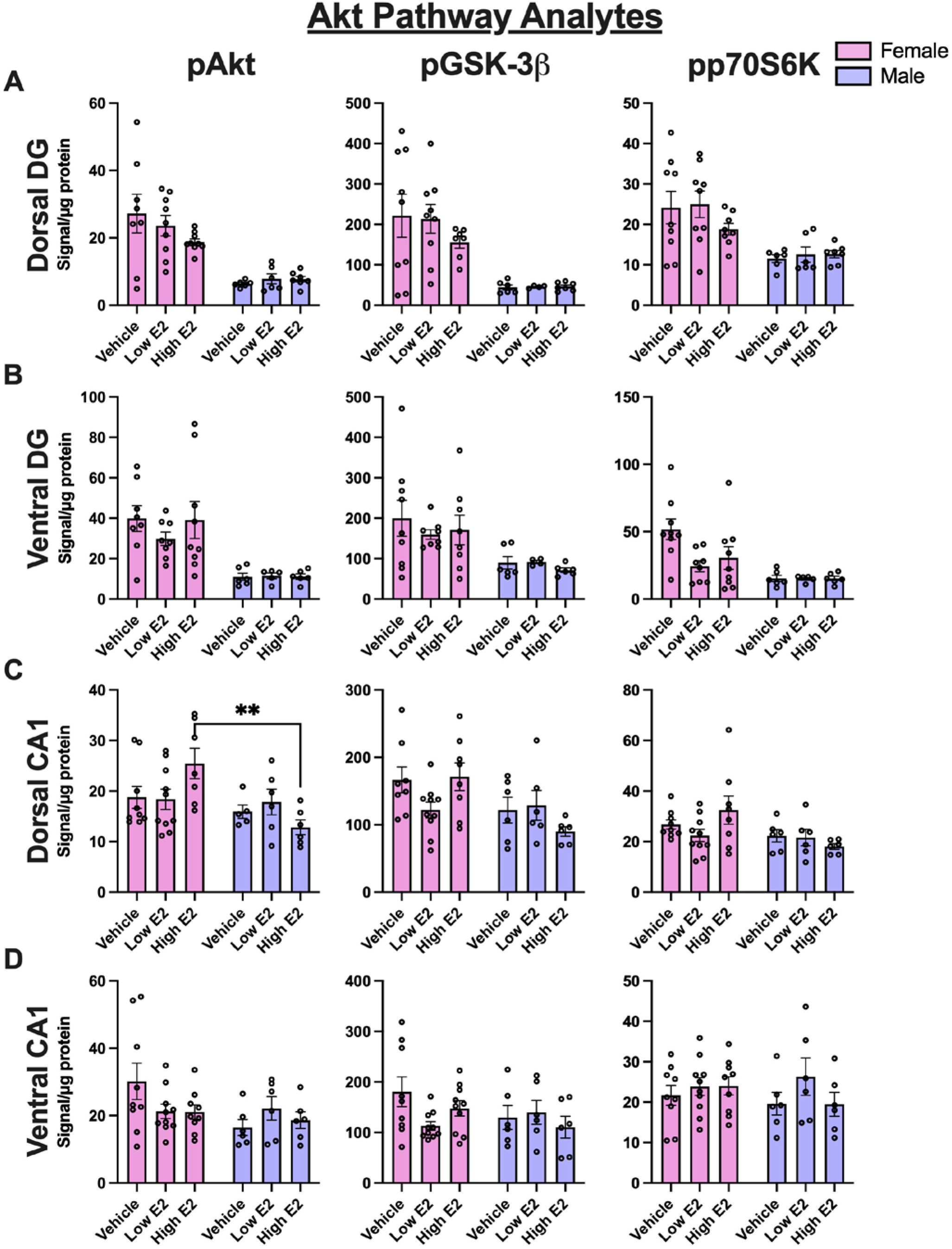
Akt phosphoprotein levels in the dentate gyrus (DG) and CA1 of female and male rats. Phosphoprotein signal normalized by amount of total protein in sample. A) Phosphoprotein levels were higher in females in all analytes in the dorsal DG (pAkt, pGSK-3β, and pp70S6K: p<0.0001). B) Phosphoprotein levels were higher in females in all analytes in the ventral DG (pAkt: p<0.0001; pGSK-3β: p=0.0004; pp70S6K: p=0.0014). C) High 17β-estradiol increased dorsal CA1 pAkt in females (p=0.05). There were no effects of sex between vehicle-treated rats. D) There were no effects of sex or treatment on Akt analytes in the ventral CA1. Error bars are ± standard error of the mean. Main effects of sex: ## p<0.01, ### p<0.001, #### p<0.0001; effects of 17β-estradiol: * p=0.05.

### Intact males and females

#### Akt pathway: Females have higher levels of Akt pathway phosphoproteins in the dorsal and ventral dentate gyrus but not in the CA1 region

Female rats had up to 400% higher levels of pAkt, pGSK-3β, and pp70S6K than males in the dorsal and ventral DG, but not in the CA1 region. Females had greater phosphoprotein signal than males for each analyte in the Akt pathway in both the dorsal (109-400%) and ventral DG (123-262%; main effects of sex: dorsal DG: pAkt – F_(1,39)_=58.51, p<0.0001, η^2^=0.262; pGSK-3β – F_(1,36)_=37.21, p<0.0001, η^2^=0.488; pp70S6K –F_(1,39)_=22.33, p<0.0001, η^2^=0.358; Figure 2A; ventral DG: pAkt – F_(1,36)_=51.22, p<0.0001, η^2^=0.581; pGSK-3β – F_(1,35)_=15.67, p=0.0004, η^2^=0.292; pp70S6K – F_(1,37)_=11.96, p=0.0014, η^2^=0.205; Figure 2B). There were no other main or interaction effects in this pathway (all ps>0.094).

High 17β-estradiol increased dorsal CA1 pAkt by 36% over vehicle treatment in females (p=0.05, Cohen’s d=0.929), but there were no other significant pairwise comparisons (interaction: F_(2,37)_=3.731, p=0.0334, η^2^=0.143; Figure 2C)). Although there were main effects of sex in the dorsal CA1 in the Akt pathway (pAkt – F_(1,37)_=7.725, p=0.0085, η^2^=0.148; pGSK-3β – F_(1,38)_=7.608, p=0.0089, η^2^=0.143; pp70S6K – F_(1,39)_=5.796, p=0.0209, η^2^=0.117; Figure 2C), *a priori* comparisons failed to show a sex differences in the vehicle conditions (all ps>0.05; Figure 2C). There were no other significant effects in the ventral CA1 (all ps>0.079; Figure 2D).

#### MAPK pathway: Males had greater pERK1/2 in the dorsal dentate gyrus than females. 17β-estradiol reduced pp38 in the ventral DG in intact females but not in intact males

Male rats had significantly greater pERK1/2 (292%) than females in dorsal DG (p=0.0004, Cohen’s d=1.389; main effect of sex: F_(1,37)_=23.87, p<0.0001, η^2^=0.372; Figure 3A). Despite the significant main effects for sex for the other analytes (pJNK – F_(1,41)_=5.832, p=0.0203, η^2^=0.117; pp38 – F_(1,42)_=38.25, p<0.0001, η^2^=0.427), there were no significant differences in pJNK or pp38 between vehicle treated males and females (ps>0.05) or other significant main or interaction effects in the dorsal DG (all ps>0.257).

**Figure 3:**
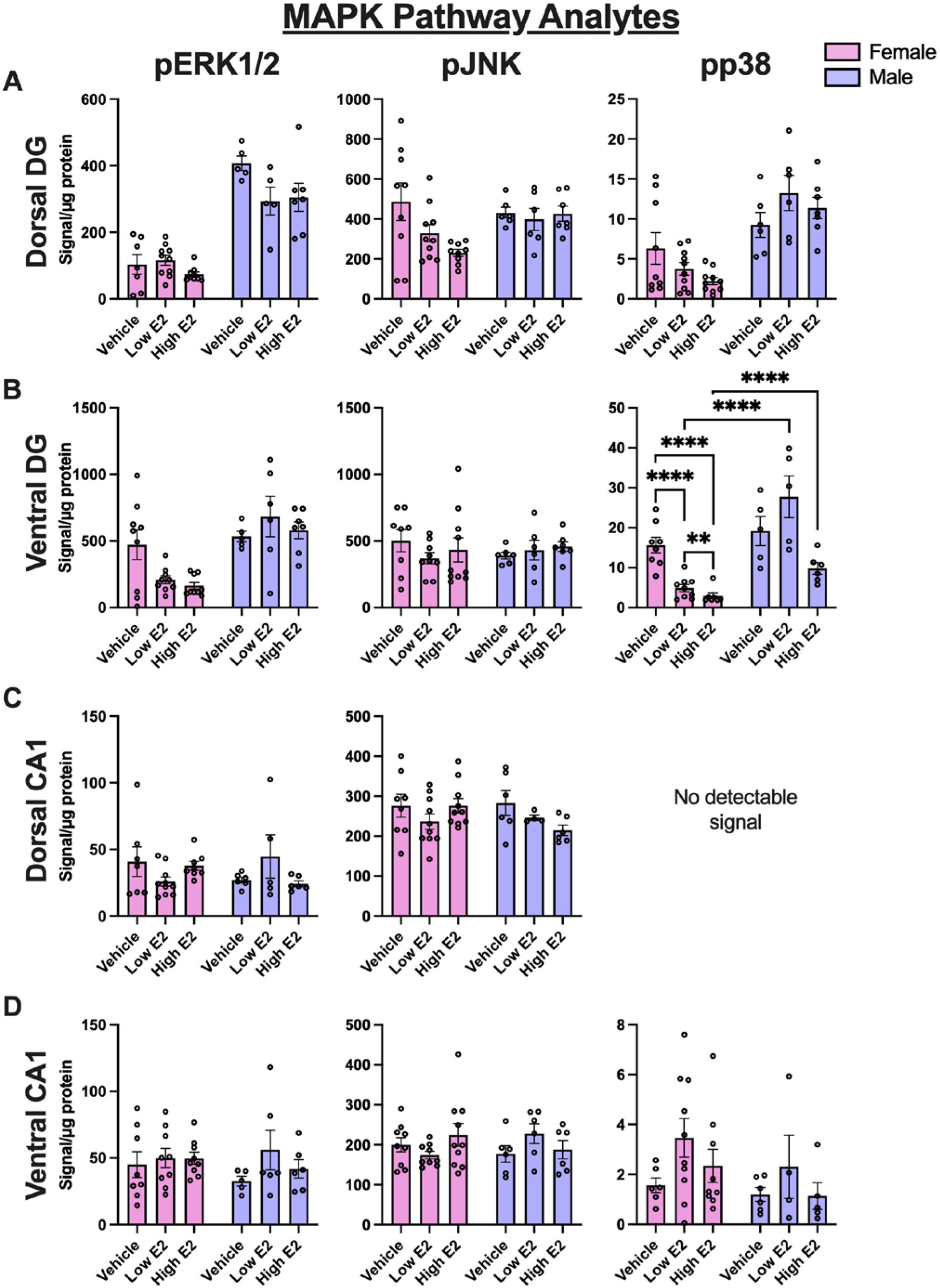
MAPK phosphoprotein levels the dentate gyrus (DG) and CA1 of in female and male rats. Phosphoprotein signal normalized by amount of total protein in sample. A) Males had higher pERK1/2 (p<0.0001) in the dorsal DG than females. There were no significant effects of sex among vehicle-treated rats in the dorsal DG for pJNK or pp38. B) In the ventral DG, there were no significant effects of sex among vehicle-treated rats. Low 17β-estradiol reduced ventral DG pp38 in females (p<0.0001), with high 17β-estradiol reducing pp38 still further (compared to vehicle, p<0.0001; compared to low 17β-estradiol p<0.01). C-D) There were no significant effects of sex or treatment on MAPK analytes in the dorsal CA1 (C) or the ventral CA1 (D). Error bars are ± standard error of the mean. Main effects of sex: #### p<0.0001; effects of 17β-estradiol: ** p<0.01, **** p<0.0001.

Both doses of 17β-estradiol decreased pp38 in the ventral DG of females (low 17β-estradiol: p<0.0001, Cohen’s d=2.260; high 17β-estradiol: p<0.0001, Cohen’s d=3.876; Figure 3B; interaction: F_(2,34)_=8.955, p=0.0008, η^2^=0.128). There were also main effects of sex on pp38 (F_(1,34)_=45.13, p<0.0001, η^2^=0.323) and 17β-estradiol (F_(2,34)_=18.54, p<0.0001, η^2^=0.266). Although there was main effect of sex for pERK1/2 (F_(1,39)_=6.067, p=0.0183, η^2^=0.133; Figure 3B), the vehicle treated males and females did not significantly differ (p>0.05). There were no significant effects on pJNK in the ventral DG (all ps>0.415).

There were no significant sex (all ps>0.145) or 17β-estradiol (all ps>0.134) effects observed for any MAPK analyte in either the dorsal or ventral CA1 in intact males or females (Figures 3C-D).

### Ovariectomized females

#### Akt pathway: 17β–Estradiol increased Akt phosphoproteins in the CA1 region of OVX rats

In ovariectomized rats, 17β-estradiol had more significant effects on cell signalling phosphoproteins in the hippocampus than in intact rats but these effects were largely restricted to the CA1 region. There were no significant effects of 17β-estradiol on Akt pathway phosphoproteins in the DG of OVX rats (all ps>0.270, Figures 4A-B). In the dorsal CA1, 17β-estradiol increased pGSK-3β by ∼34-43%, regardless of dose (43% increase with low 17β-estradiol: p=0.024, Cohen’s d=2.37; 34% increase with high 17β-estradiol: p=0.032, Cohen’s d=1.32; main effect of 17β-estradiol: F_(2,13)_=4.889, p=0.026, η^2^=0.4293; Figure 4C). However, in the ventral CA1 only the low dose of 17β-estradiol increased pAkt by 89% (p=0.042, Cohen’s d=1.397; Figure 4D) and pGSK-3β by 79% (p=0.008, Cohen’s d=1.859; Figure 4D). There were no other 17β-estradiol effects in the dorsal or ventral CA1 (all ps>0.205).

**Figure 4:**
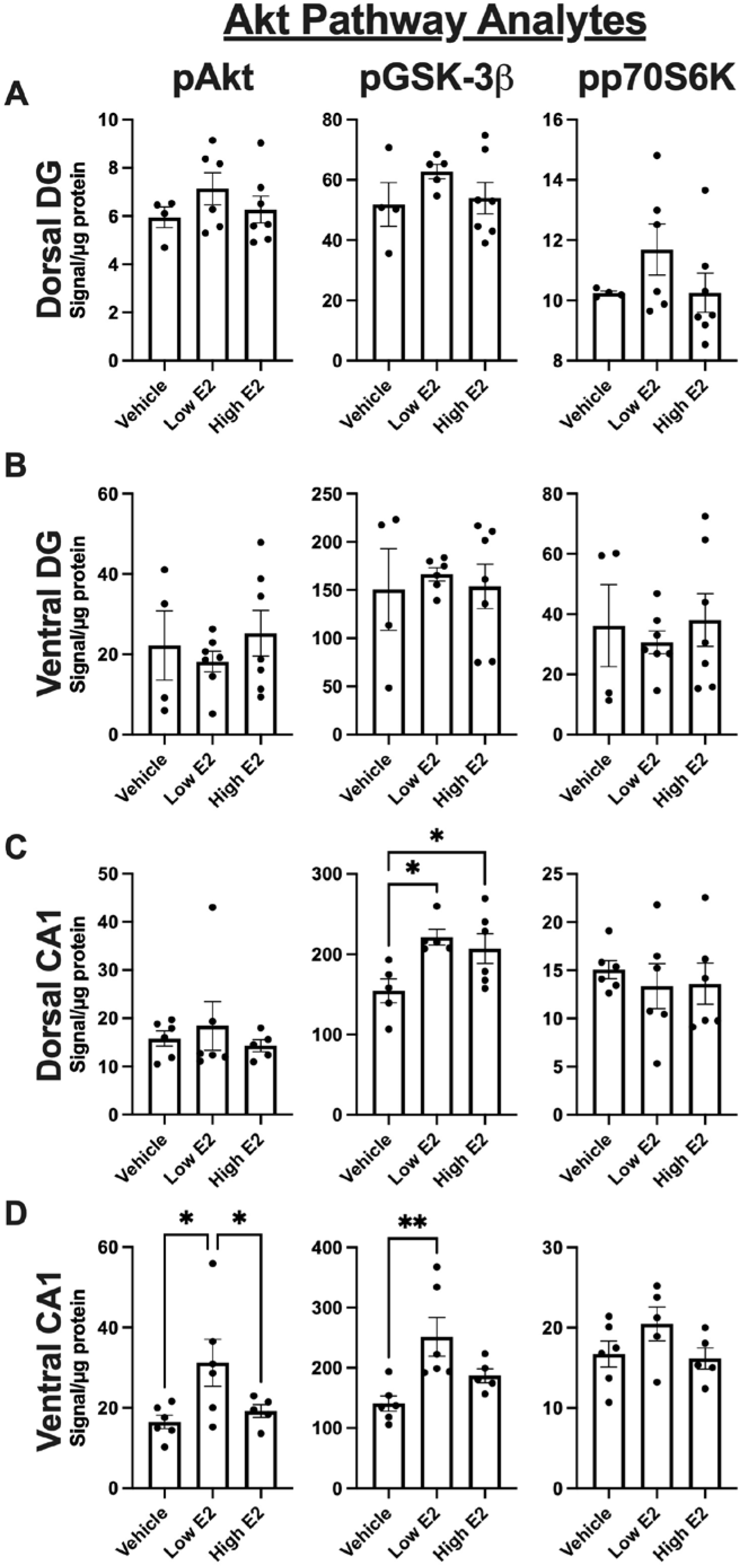
Akt phosphoprotein levels in the dentate gyrus (DG) and CA1 of ovariectomized female rats. Phosphoprotein signal normalized by amount of total protein in sample. A-B) 17β-estradiol did not affect Akt pathway phosphoprotein levels in the dorsal DG (A) or ventral DG (B) of OVX rats (ps>0.27) C) Low (p=0.0113) and high (p=0.0306) 17β-estradiol increased pGSK-3β in the dorsal CA1. D) In the ventral CA1, low 17β-estradiol increased pAkt (relative to vehicle [p=0.0142] and high 17β-estradiol [p=0.0471]) and pGSK-3β (p=0.0025). Error bars are ± standard error of the mean SEM. Effects of 17β-estradiol: * p<0.05, ** p<0.01.

#### MAPK pathway: 17β– Estradiol increases MAPK phosphoproteins in the CA1 and DG of OVX rats

17β-estradiol increased pERK1/2 and pJNK activation in the dorsal DG, (pERK1/2: low 17β-estradiol – 147% increase, p=0.031, Cohen’s d=1.356; pJNK: low 17β-estradiol – 57% increase, p=0.028, Cohen’s d=1.401, main effect of 17β-estradiol: F*_(2, 6.337)_=5.56, p=0.0403; Figure 5A). There were no other significant effects of 17β-estradiol in the dorsal or ventral DG (all p>0.303, Figure 5B).

**Figure 5:** MAPK phosphoprotein levels in the dentate gyrus (DG) and CA1 of ovariectomized female rats. Phosphoprotein signal normalized by amount of total protein in sample. A) Low 17β-estradiol increased pERK (p=0.031) and pJNK (p=0.028) in the dorsal DG of OVX rats. B) 17β-estradiol did not affect MAPK pathway phosphoprotein levels in the ventral DG of OVX rats (ps>0.303). C) Low (p=0.017) and high (p=0.009) 17β-estradiol increased pJNK in the dorsal CA1 of OVX rats. D) In the ventral CA1, low 17β-estradiol increased pERK1/2 (relative to vehicle [p<0.01] and high 17β-estradiol [p<0.05]) and pJNK (relative to vehicle and high 17β-estradiol, ps<0.05). Error bars are ± standard error of the mean. Effects of 17β-estradiol: * p<0.05, ** p<0.01.

In the dorsal CA1, both 17β-estradiol doses increased pJNK by 27% (low 17β-estradiol: p=0.017, Cohen’s d=1.56; high 17β-estradiol: p=0.009, d=1.89; main effect of 17β-estradiol: F_(2,15)_=6.288, p=0.0104, η^2^=0.456; Figure 5C). There were no other significant effects on MAPK phosphoproteins in the dorsal CA1.

In the ventral CA1, low 17β-estradiol, but not high 17β-estradiol, increased pERK1/2 by 114% and pJNK by 58% in OVX rats (pERK1/2 – p=0.003, Cohen’s d=2.434, main effect of 17β-estradiol: F_(2,15)_=8.731, p=0.0031, η^2^=0.538; pJNK – p=0.027, Cohen’s d=1.537, main effect of 17β-estradiol: F_(2,14)_=5.64, p=0.0160, η^2^=0.446; Figure 5D). There were no other significant effects of 17β-estradiol of any MAPK phosphoproteins in the ventral CA1 (all ps>0.22).

## Discussion

Past characterization of the rapid effects of estrogens on cell signaling cascades has predominantly focused on the CA1 of the dorsal hippocampus or the dorsal hippocampus as a whole and has rarely investigated sex differences. Here, we found sex differences in phosphoprotein levels in the dorsal and ventral DG and dorsal CA1 in MAPK and Akt cascades. 17β-estradiol had more dramatic effects in OVX females compared to intact females and males on MAPK and Akt pathways. In intact rats, 17β-estradiol significantly decreased MAPK phosphoproteins in females but a low dose of 17β-estradiol increased pp38 in males in the ventral DG. Dose-dependent effects of 17β-estradiol were observed mainly in the CA1 in OVX female mice in both Akt and MAPK pathways. Collectively, these findings are the first characterization of sex differences in MAPK and Akt phosphoprotein levels across the anatomical longitudinal axis of the hippocampus in the DG and CA1. These findings illustrate that both sex and 17β-estradiol differentially influence phosphoprotein levels in region-specific ways in the hippocampus.

### Sex differences in MAPK and Akt cell signalling pathways in the DG

Consistent with investigations of other brain regions, we observed sex differences in cell signaling pathways in the dorsal and ventral DG, mostly in the Akt pathway (see Table 1 for summary). Females had higher Akt pathway phosphoproteins across the anatomical longitudinal axis and between subregions of the hippocampus than males. On the other hand, males only had higher pERK1/2 than females in the dorsal DG. These findings are partially consistent to past work in other areas of the brain, as female rats had higher baseline levels of pERK1/2, pAkt, and pGSK-3β in the frontal cortex (prelimbic and infralimbic cortices), nucleus accumbens, and rostral caudate putamen than males (26). Those findings, at least the Akt pathway proteins, mirrors our findings in the hippocampus. In male mice, pERK1/2 expression in the AVPV was higher, irrespective of gonad status, compared to females (28), similar to our findings in the dorsal DG. Regrettably, while much has been made of sex differences in MAPK and Akt pathway proteins in skeletal muscle, heart, and other peripheral tissues (e.g. (46,47)), these are few investigations into sex differences in MAPK or Akt signaling in the brain. Our findings suggest sex differences, favouring females, in the Akt signalling pathways across the dorsal and ventral axis of the DG of the hippocampus and with males showing greater activation of ERK1/2 in the dorsal DG.

It is intriguing that females had higher levels of Akt phosphoproteins than males in the DG, which adds to the literature showing greater Akt phosphoproteins in the brain (26,48) and periphery (46) of females compared to males. The Akt pathway provides a mechanism through which extrinsic factors such as neurotrophic factors can activate transcription factors to affect the regulation of neural stem cells and neurogenesis in the DG (49). Recent evidence suggests that Akt pathway activation can enhance ischemia-induced neurogenesis and cell migration in males (50) and inactivation can attenuate the exercise-induced neurogenesis in the DG in male mice (51). The fact that the DG was the region that showed sex differences suggests that this may be involved in sex differences in neurogenesis that exist in maturation timelines (52) but also in neurogenic response to androgens (53) and stress exposure (54). Voluntary running increases neurogenesis in the DG of females more so than in males (55,56), suggesting another possible mechanism for this sex difference in Akt phosphoproteins. Collectively these findings suggest that one function of the increased Akt phosphoproteins may be differences in DG neurogenesis in response to certain neurotropic factors in females compared to males. Clearly more work needs to be done to determine to determine the sex specific roles of Akt in the hippocampus.

Understanding the functional significance of these differences is perhaps more important than understanding from whence they arise. Convergent sex differences – those where the underlying cellular mechanisms differ but the behavioural outcome is similar between the sexes – have been observed now in a number of brain regions and behaviours, including spatial tasks, social learning, eye-blink learning, fear behaviors, and synaptic potentiation (57–60). Finally, these data may have implications with regards to latent sex differences; that is, sex differences that only become apparent following environmental or pharmacological interventions and/or aging/disease. For example, females are resistant to the behavioural effects of the neonatal ventral hippocampal lesion model of schizophrenia, potentially resulting from greater MEK-ERK pathway activity in the prefrontal cortex and striatum (26). Perhaps greater Akt pathway phosphoproteins present in the DG of females would make them more resilient to disruptions of this pathway. We suggest that when assessing behaviour and its underlying mechanisms that researchers pay close attention to the potential sex differences in those underlying mechanisms, especially in the case of cell signalling cascades, even when no change in behaviour is present between the sexes.

### 17β-estradiol on hippocampal cell signalling pathways in OVX rats

We found more dramatic effects of 17β-estradiol on cell signalling activation in the hippocampus of OVX rats compared to intact rats (see Table 2 for summary). Estradiol rapidly increased MAPK and Akt pathway phosphoproteins dependent on dose and longitudinal axis within the CA1 region of OVX rats. It is perhaps not surprising that the CA1 region was particularly sensitive to activation of cell signalling proteins in response to 17β-estradiol as others have found that the dorsal CA1 region shows rapid upregulation of spine density and memory consolidation (27,33,34,36,61–64). Our results are consistent with Koss et al. (2018) who found that whole dorsal hippocampi of OVX female mice, but not gonadally-intact males had increased pAkt and pERK2 (p42 isoform) following acute 17β-estradiol treatment. Although we did not see statistically significant effects in the dorsal CA1 or ventral DG, the direction of the means favours an increase in pERK1/2 with 17β-estradiol. It is important to note here that Koss and colleagues (27) saw differences in pERK only in pERK2 and not pERK1 whereas our analysis did not discriminate between isoforms. In addition to the hippocampus, estradiol rapidly increases pERK1/2 in the medial preoptic area and AVPV (28). Further, effects of 17β-estradiol on pAkt were present only in the ventral CA1 and not in the dorsal CA1, which is interesting given the functional implications of these regions. Future research should consider the anatomical longitudinal axis, as findings can differ based on this and may be important for therapeutic outcomes.

As alluded to above, 17β-estradiol treatment increased pERK1/2 in the DG and ventral CA1 of OVX rats but not in gonadally intact male and female rats. Consistent with our findings, Koss et al (27) found that 17β-estradiol increased pERK2 in OVX female mice, and not in intact males. Much of the previous work on the rapid effects of 17β-estradiol has focused on comparing to an OVX group. However, except for pAkt in the dorsal CA1 and pp38 in the ventral DG, the effects of 17β-estradiol treatment in this study were only present in OVX rats and not intact rats. It is possible that more effects would be seen at a different (earlier) timepoint than 30 minutes (discussed below). Another possible reason is that the intracellular response to estrogens is already at ceiling in gonadally-intact female rats. While not directly compared, phosphoprotein levels (e.g. Akt analytes in the dorsal DG) in OVX rats were lower than in gonadally-intact females. Further supporting this is the inverted U-shaped dose response in OVX rats in which low 17β-estradiol increased phosphoproteins whereas high 17β-estradiol largely did not. This is frequently seen in sex steroid research (9,65) and likely results from higher doses activating off target effects. As such, treatment of gonadally-intact female mice with 17β-estradiol may have resulted in fewer effects on MAPK and Akt analytes due to the ongoing presence of gonad-derived hormones. This has substantial implications for the conclusions and translatability of preclinical 17β-estradiol research. For example, if 17β-estradiol treatment leads to an improvement in a cognitive task in OVX rodents in a MAPK-or Akt-dependent manner, it should not be assumed that the same improvements would be seen in gonadally-intact rodents.

### Differences across hippocampal subregions and along the longitudinal axis

Estrogens, in particular 17β-estradiol, affect neuroplasticity in a subregion-dependant manner in the hippocampus. For example, 17β-estradiol increases apical dendritic spine density in the dorsal CA1 (21,25,31,61–64) but not in the DG (21,31). However, 17β-estradiol rapidly increases cell proliferation in the dorsal and ventral DG of OVX rats (6,30), with the effects on males or gonadally-intact females to be determined. Inhibition of ERK signaling blocks the increased dendritic spine densities in dorsal CA1 and medial prefrontal cortex following intrahippocampal 17β-estradiol treatment (64). Given our findings that low 17β-estradiol increases pERK, pJNK, and pGSK-3β in dorsal DG, perhaps the rapid effects of 17β-estradiol on cell proliferation in the DG (6) involves activation of these phosphoproteins. As such, investigations into the intracellular mechanisms driving rapid increases in cell proliferation in the DG are required.

It remains possible that the other subregions of the hippocampus drive effects seen in previous studies that were absent in ours. Here, we focused on the CA1 and DG as many of the established rapid effects of estrogens occur within these regions (e.g. (6,30,61–64); for recent reviews, see (3,9)). As many previous studies had less regional specificity, perhaps the effects, such as increases in pERK in the dorsal hippocampus response to 17β-estradiol, are driven by hippocampal regions other than the CA1 and DG, such as the CA3 region.

### Estrous cycle

In the present study, we did not observe any statistical effects of estrous cycle, although our study was not designed to specifically examine estrous cycle changes. There is a paucity of research making comparisons across the estrous cycle with regards to cell signaling cascades in the brain. One study found that transcription of MAPK and, potentially, Akt cascade phosphoprotein mRNA in the medial prefrontal cortex of rats was upregulated in proestrus (66). In the ventral hippocampus, chromatin dynamics and resulting MAPK pathway protein gene expression vary across the estrous cycle and between male and female mice (67). In addition to changes in dendritic spine densities in the rat hippocampus across the estrous cycle (68,69), changes in total hippocampal volume across the mouse estrous cycle have been observed, with high estrogen phases associated with greater hippocampal volume (70). Similar changes have been found across the human menstrual cycle (71,72). With so many neuroplastic changes occurring with natural fluctuations of estrogens, further investigations into the effects of the estrous cycle on estrogen-responsive kinase pathways are needed.

### Limitations and Future Directions

One limitation is that we only examined a 30 min window after 17β-estradiol exposure. We selected a 30 min delay in the present study as others have noted changes to hippocampal physiology and behaviour have been observed at this timepoint (6,30,64,73) following 17β-estradiol or shortly thereafter (e.g 40 min (61–63)). However, other studies (using intrahippocampal infusion) have found increases in many MAPK and Akt phosphoproteins in the hippocampus as early as 5-10 min following 17β-estradiol (33–36). Due to the transient nature of phosphorylation in cell signalling cascades, it remains possible that the effects of treatment occurred at an earlier timepoint than in our study. Furthermore, it is possible that both sex and/or gonadal status may affect the rate at which these rapid effects of 17β-estradiol occur and should be investigated further.

The present data were normalized to total protein in sample (as is done in regularly in the literature: (39,43–45), for example). One possible limitation is that phosphorylated proteins were not normalised to the specific protein under investigation (e.g. pERK1/2 normalized to total ERK1/2). It remains possible that there are sex differences in and estradiol effects on total protein levels that could confound the interpretation of the present results. Genomic synthesis of novel MAPK, Akt, or other proteins triggered by 17β-estradiol is unlikely due to the time of tissue collection (30 min). It remains possible, however, that 17β-estradiol could rapidly increase the total protein levels via non-genomic mechanisms, such as the translation of mRNA transcripts already in the cell. Additionally, it is through the phosphorylation of these proteins that functional outcomes occur (e.g. DNA transcription, protein translation, actin polymerization, etc.), thus obtaining the functional unit of these pathways.

The electrochemiluminescent assay is highly sensitive and selective, provides quantitative data for analyses, and requires relatively little tissue for analysis compared to similar assays (41,42). Other cell signaling cascades or proteins, such as the eukaryotic translation initiation factor 4E-binding protein 1, a translation repressor protein downstream of many Akt and MAPK phosphoproteins including Akt, ERK1/2, and p38 (74) would be important proteins to examine in the future. Estradiol only had significant effects on phosphoprotein level in the ventral DG where pp38 was decreased in female rats only. This reduction could result from inactivation by upstream kinases or as a compensatory mechanism following transient activation (75), which would be important to examine in the future. In addition, in the present study we focussed on the CA1 and DG as they are rapidly affected by estrogens (9). However, the CA3 is an additional region of interest, given sex and estradiol differences in morphology and response to stress in this region (20,68,76).

### Conclusions

Sex differences in MAPK and Akt cell signaling phosphoproteins exist in both the dorsal and ventral DG in rats. Estradiol treatment increased MAPK and Akt signaling in the CA1 and dorsal DG in OVX rats with far fewer effects in intact females or males. These results shed light on the possible underlying mechanisms of sex differences in hippocampal function in response to 17β-estradiol that affect MAPK and Akt kinase pathways. They further show the heterogeneity of estrogen and sex effects between hippocampal subregions and along the dorsal/ventral axis. With these subregions serving often distinct purposes, and the functional dissociation between the dorsal and ventral hippocampus, the present results provide possible mechanisms through which these differences may arise. Furthermore, with many neurological disorders having profound effects on the hippocampus, these sex, hormonal status, and regional differences provide valuable insights not only into the potential underlying mechanisms, but also into therapeutic possibilities.

## Acknowledgements

The authors would like to thank Stephanie Lieblich for her assistance with surgeries, Dr. Rand Eid for her guidance and assistance with the electrochemiluminescence assays, Muna Ibrahim and Rebecca Rechlin for estrous cycle determination, and the staff at the Centre for Disease Modeling at the University of British Columbia for their ongoing care for and kindness towards the animals in that facility.

## Statement of Ethics

All experiments were conducted in accordance with the ethical guidelines set by the Canada Council for Animal Care and were approved by the University of British Columbia Animal Care Committee. All efforts were made to reduce the number and the suffering of animals.

## Author Contributions

PASS – conception and design of the work; data acquisition, analysis, and interpretation; writing and revising

TAP -conception and design of the work; data acquisition, analysis, and interpretation; writing and revising

LAMG -conception and design of the work; data acquisition, analysis, and interpretation; writing and revising

## Data Availability

All data is available upon request to corresponding author.

